# The 3D organisation of mitochondria in primate photoreceptors

**DOI:** 10.1101/2021.06.03.446914

**Authors:** Matthew J. Hayes, Dhani Tracey-White, Jaimie Hoh Kam, Michael B. Powner, Glen Jeffery

**Affiliations:** University College London Institute of Ophthalmology 15-43 Bath Street London EC1V 9EL; City, University of London, Northampton Square, London EC1V 0HB

**Keywords:** Mitochondrion, mitochondria, photoreceptor, rod, cone, mitophagy, stress, MDV

## Abstract

Vertebrate photoreceptors contain large numbers of closely-packed mitochondria which sustain the high metabolic demands of these cells. These mitochondria populations are dynamic and undergo fusion and fission events. This activity serves to maintain the population in a healthy state. In the event of mitochondrial damage, sub-domains, or indeed whole mitochondria, can be degraded and population homeostasis achieved. If this process is overwhelmed cell death may result. Death of photoreceptors contributes to loss of vision in aging individuals and is associated with many eye diseases. In this study we used serial block face scanning electron microscopy of adult (*Macaca fascicularis*) retinae to examine the 3D structure of mitochondria in rod and cone photoreceptors. Healthy-looking photoreceptors contain mitochondria with a range of shapes which are associated with different regions of the cell. In some photoreceptors we observe mitochondrial swelling and other changes we associate with stress or degeneration of the cell. In both rods and cones we identify elongated domains of mitochondria with densely-packed normal cristae associated with photoreceptor ciliary rootlet bundles. We observe mitochondrial fission and mitochondrion fragments localised to these domains. Swollen mitochondria with few intact cristae are located towards the periphery of the photoreceptor inner-segment in rods, whilst they are found throughout the cell in cones. Swollen mitochondria exhibit sites on the mitochondrial inner membrane which have undergone complex invagination resulting in membranous, electron-dense aggregates. Membrane contact occurs between the mitochondrion and the photoreceptor plasma membrane in the vicinity of these aggregates, and a series of subsequent membrane fusions results in expulsion of the mitochondrial aggregate from the photoreceptor. These events are primarily associated with rods, likely reflecting the ageing mechanism in primates where many rods die but cones do not. Possible consequences of this atypical mitochondrial degradation are discussed.

## Introduction

Photoreceptor neuron degeneration is a natural part of ageing and disease, causing blindness in approximately 1 in 2000 people globally. More than half of such cases are a result of mutations in around 200 genes, including those responsible for retinitis pigmentosa and Leber’s hereditary optic neuropathy^1^. Degenerative changes also occur in diabetic retinopathy and age-related macular degeneration (AMD) due to a confluence of multifactorial genetic and environmental factors ^2–4^. In all these conditions mitochondrial dysfunction has been identified ^5–7^.

Rod photoreceptors represent most of the energy consumption of the retina ^8,9^. They require ATP for the transport of ions during the depolarisation and hyperpolarisation currents of phototransduction (particularly the Na+/K+ ATPase ^10,11^ during the dark current), and for lipid and protein synthesis that is required for outer segment renewal as 10% of the distal tip of the outer segment is replaced every day. More than 80% of glucose taken up by the photoreceptor is used for anaerobic glycolysis ^12^, but a proportion of ATP is provided by mitochondria ^9^. Oxygen consumption measurements suggest that the photoreceptor mitochondria operate close to capacity ^13^. The mitochondria are also vital in the compartmentation of cell metabolism and regulation of programmed cell death ^14^.

Photoreceptors are highly polarised, terminally differentiated cells and in most vertebrate species mitochondria are predominantly packed into a region of the inner segment called the *ellipsoid*.

Photoreceptors are bathed in incident radiation and exposed to reactive oxygen and nitrogen species (RONS) generated by these mitochondria and by the ionising effects of light on photosensitizing molecules such as retinoids. Oxidation of abundant long-chain polyunsaturated fatty acids (PUFAs), produce highly-reactive electrophilic intermediates that generate free-radicals. Hence, photoreceptor mitochondria accumulate damaging modifications through thermal, photoreaction and redox mechanisms ^15,16^. Mitochondria may then signal cell death by a number of programmed mechanisms including apoptosis ^17^. In primates, including humans, up to 30% of rods are lost over life, but there is little evidence for cell death in the cone population, although these suffer functional decline ^18^

Mitochondrial populations in cells are heterogeneous, their morphology changing in response to damage, stress or metabolic demand. Mitochondrial biogenesis occurs as a result of fission ^14^. By growing new mitochondria from the best elements of existing ones, the cell avoids accumulating errors. Homeostasis occurs by highly-regulated interplay between fission, fusion and autophagy ^14,19–26^. In mammalian cells, mitochondrial fusion is performed by mitofusin-1, mitofusin-2 (MFN-1/2) and optic atrophy-1 (OPA1). By sharing and mixing their contents dysfunctional mitochondria are thought to mitigate localised damage ^27^. Here we generate 3D reconstructions from serial sections of mitochondria in healthy and apparently stressed rods and cones to see if there are intracellular, localised variations in morphology that are likely to reflect dynamic homeostasis.

In primate photoreceptors a ciliary rootlet extends from the pair of basal bodies at the base of the cilium, along the length of the cell, around the nucleus and as far as the synaptic termini. In the transmission electron microscope, it appears striated in longitudinal section, the banding representing overlapping components of the intermediate filament rootletin ^28^. In mutant mice deficient for rootletin, the outer segment forms as normal, but soon degenerates, suggesting that the rootlet is involved in maintenance or stabilisation of the outer-segment ^29^. The rootlet is inherently ‘non-directional’ being composed of symmetrically arranged elements having no positive or negative end, it is thought unlikely to be used for directional trafficking of cell components.

We wanted to see if mitochondria associate with the photoreceptor rootlet in this old-world primate as has been seen in other species^3031^. We show that a proportion, those densely furnished with cristae, and exhibiting a narrow diameter, do associate with the rootlet. We identify mitochondrial fission events localised to this region. We also describe a process by which membranous, electron-dense domains of the inner mitochondrial membrane in swollen mitochondria are expelled from photoreceptors. These results suggest photoreceptor mitochondria homeostasis is a cycle of fission and piecemeal degradation associated with the rootlet and expulsion through plasma-membrane respectively.

## Methods

### Animals: primates

Ocular tissues were acquired from a large long established colony of Macaca fascicularis maintained by Public Health England regulated under local and U.K. Home Office regulation. The primary purpose of animal usage was different from the aims of this study and eyes were only retrieved after death. Primates were of Mauritian origin and have a smaller gene pool than those from Indonesia and age more rapidly. Their normal maximum lifespan is around 18 years in this colony. In all experiments retinal tissue from at least five individuals between 12 and 18 years of age was examined.

### Conventional Transmission electron microscopy

All animals were heathy and were sacrificed between 9-11am. Animals were sedated with 200mg /kg of Ketamine I/M and then terminally anaesthetised with 2.5ml of Nembutal (50mg sodium pentobarbital per mg) delivered to the heart. Eyes were removed at the point of death and were fixed overnight in cold Karnovsky’s fixative (2% paraformaldehyde, 2.5% glutaraldehyde in 0.08M cacodylate buffer). Small portions of the perifoveal retina were dissected. They were washed 3 times in phosphate buffer and osmicated with 1% osmium tetroxide in ddH_2_O for 1 hour. Samples were then washed 3 ⨯ 10 minutes in ddH_2_O and dehydrated with a series of ethanol dilutions: 30%, 50%, 70%, 90%, 3 ⨯ 100% and 2 ⨯ propylene oxide (at least 20 minutes in each). They were infiltrated with 50:50 propylene oxide:araldite resin overnight and with several changes of 100% resin the next day. Blocks were cured at 60°C overnight. Sectioning was performed using a Leica Ultracut UCT microtome. Sections were counter-stained with Reynold’s lead citrate and were viewed on a JEOL 1400+ TEM (JEOLUSA MA, USA).

### Serial block face scanning electron microscopy SBF-SEM (after Walton 1979^32^)

Specimens were fixed in Karnovsky’s EM fixative (as above) for 30 mins. They were washed 3 times with phosphate buffer. Samples were incubated for 2 hours in 2% osmium tetroxide/1.5% ferricyanide and washed 3 ⨯5 mins in water. The samples were placed in 1% thiocarbohydrazide solution for 10 minutes and then washed 3 ⨯5 mins in water. A second phase of osmication was performed (2% osmium tetroxide (no ferricyanide) for 30mins) and the tissue washed again in water. The samples were then placed in aqueous 2% uranyl acetate overnight at 4 degrees.

The following day samples were placed in freshly made Walton’s lead aspartate (pH5.5) for 30 mins in a 60°C oven and subsequently washed 3 × 5 mins in water. Samples were then dehydrated with ethanol (30%, 50%, 70%, 90%) to 100% x 3 then in acetone 2 × 20 mins. Samples were infiltrated with Durcapan resin 25% plus acetone, then 50%, 75% (2 hours each) and left overnight in 100% resin. The following day the resin was exchanged, and the blocks hardened in an oven for 48 hours. The samples were examined on a Zeiss Sigma VP SEM fitted with a Gatan 3View system.

### Quantitation of mitochondrial distribution

The distance of the mitochondria from the center of the photoreceptor and the diameters of these mitochondria in normal and degenerate photoreceptors was measured from individual transverse transmission electron micrographs using ImageJ. Results are from 4 animals and at least 5 cells in each case. Statistical differences between normal and degenerate and rootlet-associated and non-associated mitochondria were calculated by two-tailed Mann-Whitney U Tests. *** implies statistical significance > than P = 0.0001. Un-labelled pairwise comparisons failed to reach significance of P = 0.01 and were considered to be insignificant.

## Results

### 3D structure of healthy cones

In TEM sections of cone photoreceptors mitochondria appear as closely-packed oval structures. Examination of sections at different orientations gives little additional insight into their arrangement and structure. We performed serial block face scanning electron microscopy (SBF-SEM) to allow us to identify individual mitochondria across several hundred such sections and thus reveal their 3D structure and position in the photoreceptor.

Figure 1 shows a 3D reconstruction of every mitochondrion in a cone in the peripheral retina of a 14 year-old animal generated by manual segmentation from an SBF-SEM stack. The ellipsoid contains a densely-packed array of mostly elongated mitochondria arranged broadly parallel to the long axis of the photoreceptor. The cell contained 498 individual mitochondria. The apical mitochondria terminated in narrow tips whilst those that terminated distally were swollen. In this cell we identified few mitochondrial fragments; those present were mostly apical.

**Figure 1.**
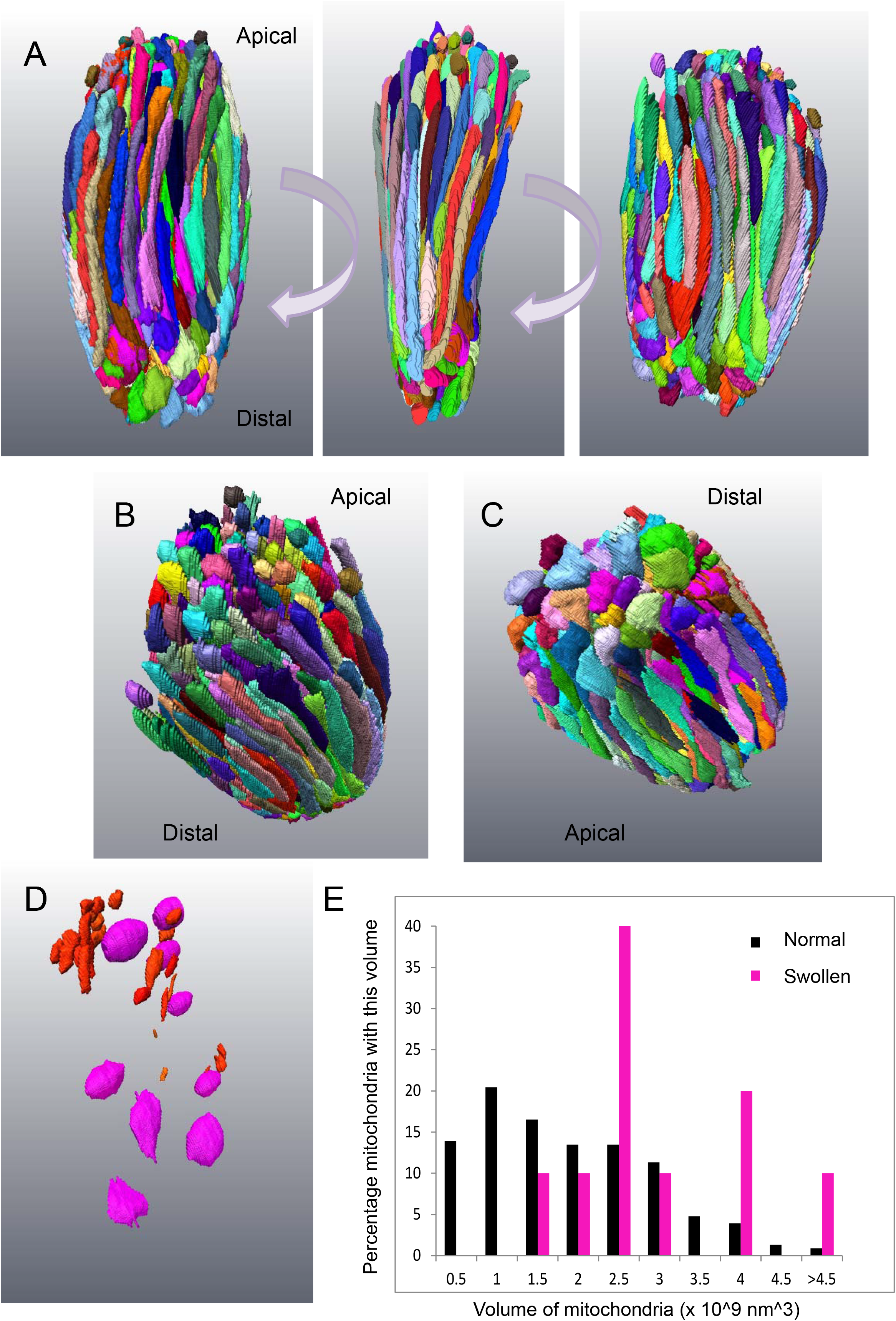
3D rendering of complete segmentation of every mitochondrion in a small, normal cone. A: Three panels represent 90 degree rotations of the ellipsoid of a cone cell showing all the mitochondria. The mitochondria show a range of profiles, though most are long and thin. B: Mitochondria at the apical end of the ellipsoid terminate with narrow tips. There were also numerous mitochondrial fragments. C: Those at the distal end were swollen at their tip. D: Mitochondrial fragments (red) are mostly localised to the apical domain of the ellipsoid. Spheroidal, swollen mitochondria (magenta) are found throughout. E: Histogram showing the range of mitochondrial volumes in this ellipsoid. The spheroidal mitochondria (magenta) have a greater average volume.

Dispersed throughout the cell were a small number of spheroidal mitochondria which had an electron lucent matrix and few cristae. The volume of these mitochondria was found to be greater than those of the denser, elongated mitochondria; indicating they were swollen as well as being of an altered shape (Fig. 2D-G).

**Figure 2.**
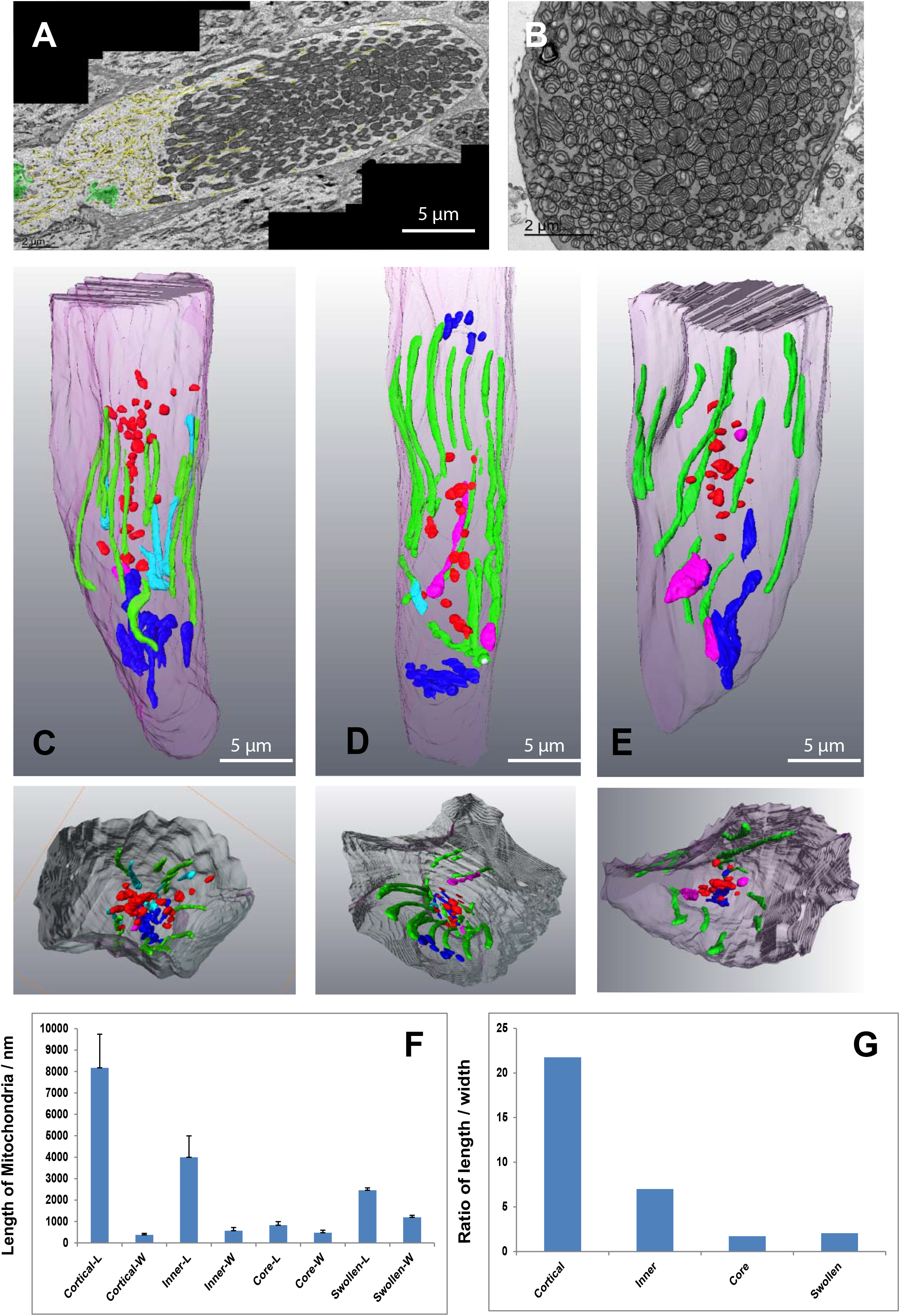
In some ‘normal’ cones different mitochondrial shapes are associated with different parts of the photoreceptor. A: A montage of several TEM micrographs of the longitudinal section of a healthy-looking cone. The ER is high-lighted in yellow, the Golgi apparatuses in green. B: A TEM micrograph showing a transverse section through a healthy-looking cone. Mitochondria are densely-packed in the ellipsoid. C-E: Partial 3D segmentation of 3 representative cones identifying different mitochondrial shapes in different regions of the cone ellipsoid. The smaller panels beneath show cut transverse sections through these constructions. F: Histogram showing the average lengths of mitochondria from different parts of the cone. G: Histogram showing the ratio between length and width of mitochondria from different regions of the cone. Mitochondria intimately associated with the plasma membrane of the cone (green) are elongated. Those inner mitochondria that are only partially associated with the plasma membrane (cyan) are shorter and have a more varied width. Mitochondria at the base of the cone are shorter and stubbier (dark blue). Mitochondria fragments in the core of the cone (red) are very small.

In individual TEM sections the mitochondria of large, parafoveal cones appear as closely-packed ovals, in transverse sections mostly as circles (Fig. 2A-B). 3D rendering of these from SBF-SEM stacks reveal them to have a range of morphologies (Fig. 2C-E). Mitochondria closest to the photoreceptor plasma membrane (cortical) exhibited extended membrane contact sites with the plasma membrane. These mitochondria were uniformly narrow and elongated, having a high length/width ratio. Those less closely associated with the plasma membrane (inner mitochondria) had slightly broader, shorter profiles. Those at the distal end were shorter and broader still (Fig. 2F-G).

Mitochondrial fragments were localised to the innermost core of the cell and tended to be more apical than distal (Fig. 2C-E).

### 3D structure of healthy rods

Figure 3A-B shows a 3D reconstruction of every mitochondrion in a small, healthy-looking rod in the central retina from a 12 year-old individual. The mitochondria are all narrow and elongated (Fig. 3A) and in transverse section (TEM) appear mostly circular (Fig 3D). In such sections it is possible to identify the ciliary rootlet. Some mitochondria appear to be in close contact with this structure. In the schematic the ciliary rootlet is highlighted in red and rootlet-associated mitochondria in blue.

**Figure 3.**
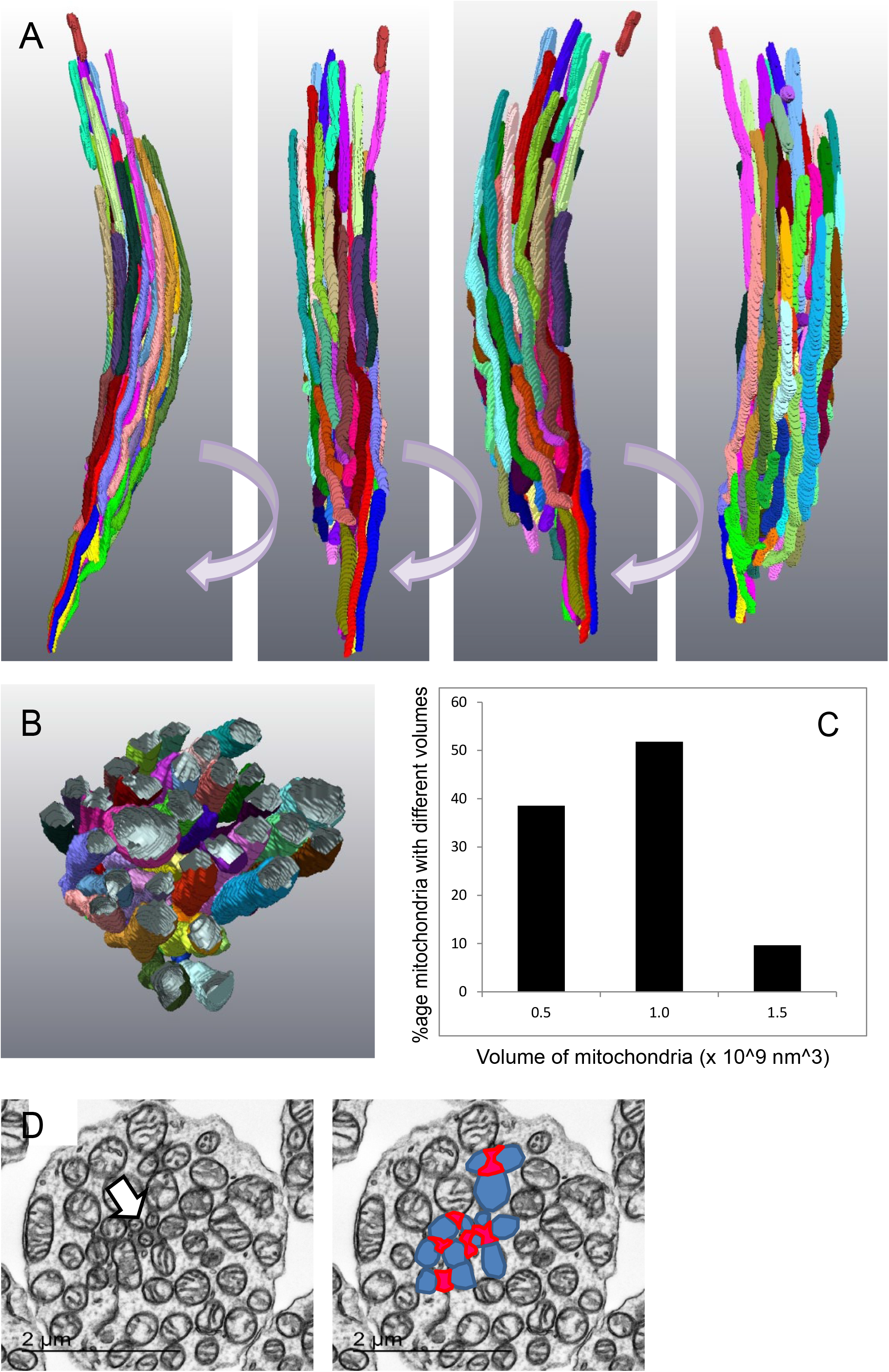
3D rendering of complete segmentation of every mitochondrion in a normal rod. A: Rotations of a 3D reconstruction of segmentation of mitochondria in a normal, healthy-looking rod. All the mitochondria are elongated and aligned along the long axis of the rod. B: Transverse section generated from the 3D reconstruction. C: Distribution of mitochondria volumes. D: Transverse section showing the position of the rootlet and its association with mitochondria (high-lighted in the next panel); ciliary rootlet in red, associated mitochondria in blue.

### Mitochondria in stressed cone photoreceptors

As described above, mitochondria in healthy-looking cones appear to have a dense matrix and are rich in cristae. In some cases we observed mitochondrial extensions along the ciliary rootlet (Fig. 4A).

**Figure 4.**
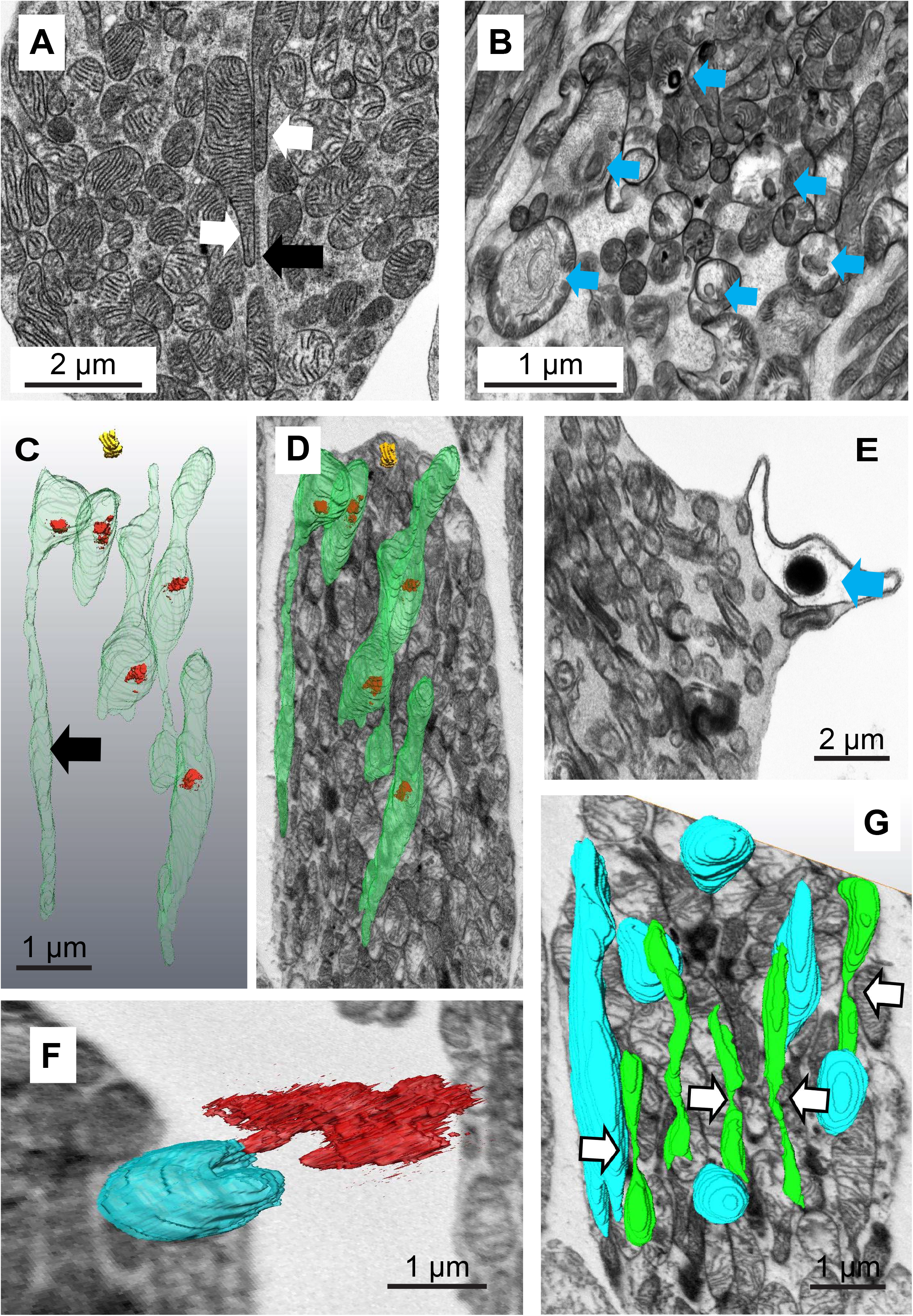
Mitochondria in stressed cones. A: Oblique section through a healthy-looking cone showing two mitochondria with extensions (white arrows) associated with the ciliary rootlet (black arrow). B: Oblique section through a stressed cone illustrating a range of mitochondria forms (mitochondria containing few cristae and electron-dense aggregates marked with a blue arrow). C: 3D reconstruction of mitochondria containing electron dense inclusions D: Super-position of this reconstruction on a single SBF-SEM section showing their position in the cell. E: A swollen mitochondrion containing an electron-dense inclusion associated with the plasma membrane. F: A 3D reconstruction from a SBF-SEM stack showing a C-shaped, plasma-membrane-associated, swollen mitochondrion which has expelled material through the plasma membrane. G: A 3D reconstruction of segmentation from a SBF-SEM stack showing normal mitochondria (blue) and narrow mitochondria (green) apparently undergoing fission (white arrows).

Generally, in cones we saw few atypical, swollen mitochondria or inclusions (consistent with their survival through ageing compared with rods); however, some regions of the adult retina exhibited cones which appeared to be stressed. In some cases these were in very close proximity to dead rods (identified as being highly condensed, with little or no outer segment and having few identifiable remaining organelles). Stressed cells were identified as having swollen or heterogeneous mitochondria but in which the outer segment, nucleus, endoplasmic reticulum and Golgi appeared normal. There was no evidence of overt apoptosis or necrosis (Fig. 4B).

A minority of mitochondria in these cells had empty regions of matrix, few linear cristae and instead contained aggregates of electron-dense deposits surrounded by what appear to be concentric or spirally arranged whorls of membrane (see arrows in Fig. 4B).

Reconstructions from segmentation of SBF-SEM stacks of cones revealed most of these aggregates to be fully internal to the bounding outer mitochondrial membrane (Fig. 4C-D). Under the sample preparation conditions used for SBF-SEM, the contents of the cristae of the mitochondria appear electron-dense and the mitochondrial inter-membrane space is also packed with dense material (though less so than the cristae). The electron-dense inclusions are associated with the mitochondrial periphery. We identify a region of electron-lucid mitochondrial inter-membrane (MIM) space in the vicinity of these inclusions suggesting that the dense components have been drawn into the whorl of membrane (Fig. 5A white arrow).

**Figure 5.**
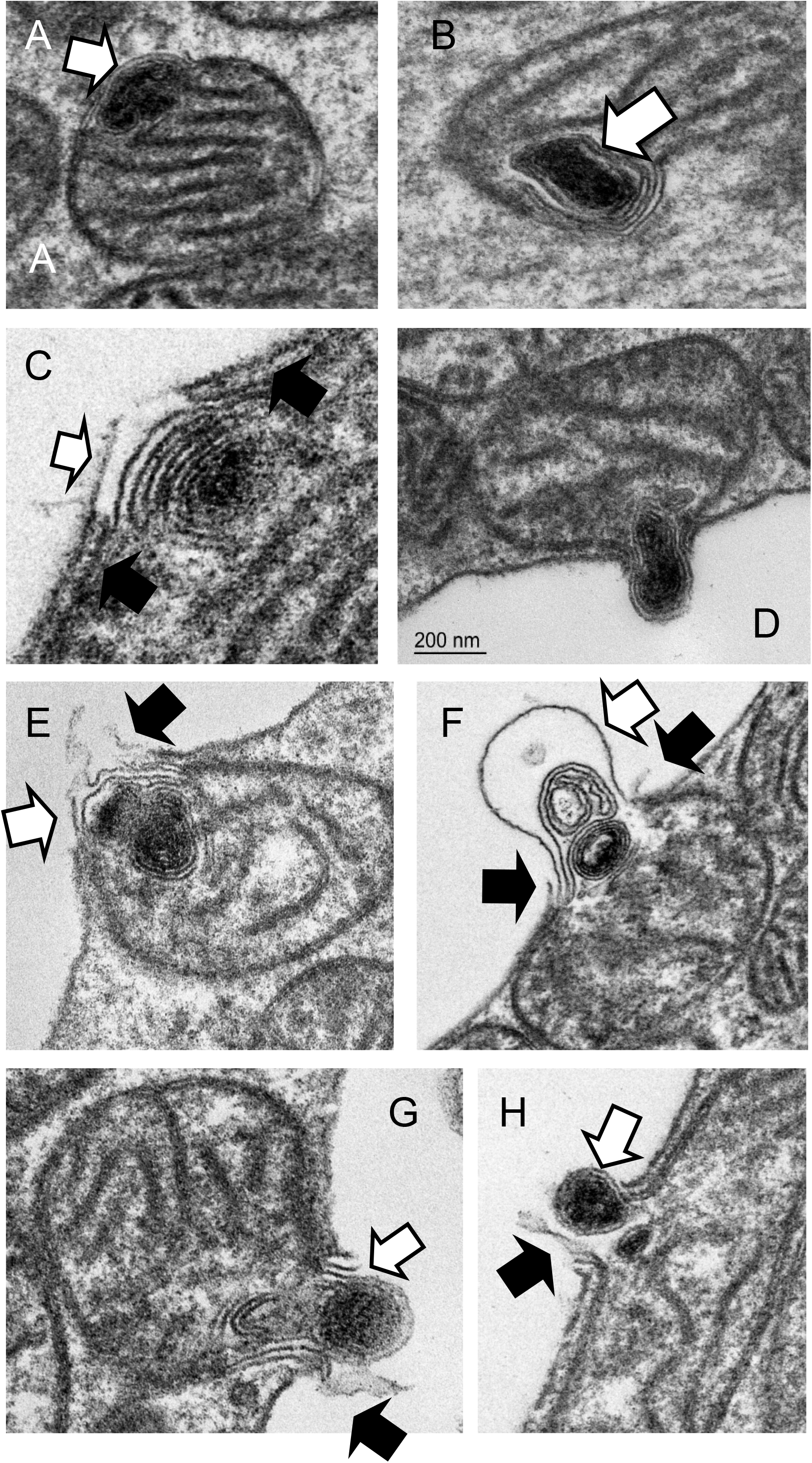
Mitochondria in stressed cones expel electron-dense, membranous contents through the plasma membrane. A-B: Electron dense membranous aggregates in mitochondria are associated with electron-lucent domains of the mitochondrion inter-membrane space (white arrows). C-H: Mitochondria engage with the plasma membrane, undergo a series of membrane fusions and eject the membranous aggregate into the inter photoreceptor space. (White arrows: swollen or ruptured mitochondrial outer membrane, black arrows: ruptured plasma membrane.

More than 90% of cones exhibited few obvious mitochondrial morphological changes; but most contained a number of mitochondria containing electron-dense material. Examination of cones from the central retina from five specimens (ages 14-18 years) suggested to us a speculative series of events leading to the mitochondria expelling the electron-dense material through the plasma membrane of the photoreceptor (Fig. 5A-H). The dense aggregate forms in the vicinity of the inter-membrane space and this region of MIM is altered (Fig. 5A-B). In cones this appears stable as we never observe aggregates free in the cytoplasm. If the mitochondrion contacts the photoreceptor plasma membrane, the outer mitochondrial membrane in this altered domain promotes partial membrane rupture or fusion (Fig. 5C-D). Subsequently, the mitochondrial membranes swell out through the rupture in the plasma membrane and are likely expelled into the inter-photoreceptor space (Fig. 4E-F, Fig. 5E-H). We presume the plasma membrane and mitochondrial inner and outer membrane reseal and the mitochondrion, now purged of its aggregate, can return to the cell population.

In cones exhibiting heterogeneous mitochondria we also observed regions of elongated, narrow mitochondria in which we could identify sites of possible mitochondrial fission (Fig. 4G).

### Mitochondria in stressed rod photoreceptors

We identified regions of apparent rod photoreceptor stress where nearby cones often appeared healthy. Stressed rods appeared slightly shrivelled but with normocytic nuclei and extended, normal-looking outer-segments (data not shown). In cross-section, they exhibited swollen mitochondria with electron lucent matrix and few cristae surrounding what appear to be a core of narrow, dense, mitochondria associated with the ciliary rootlet (Fig 6A-C). In transverse section it is apparent that these are in fact tadpole-shaped mitochondria composed of swollen domains with few cristae connected to elongated tubular extensions (Fig. 6D). The cristae of these narrow domains partially align with the repeats of the ciliary rootlet (Fig. 6E). Examination of the distance of the mitochondrion from the geographical centre of the cell and an examination of size reveal that in healthy rods rootlet-associated mitochondria tend to be in the core of the photoreceptor and are slightly narrower than those at the cortex of the cells. In stressed cells those that are associated with the rootlet exhibit a narrow profile, whilst those excluded from the rootlet are swollen (Fig. 6F).

**Figure 6.**
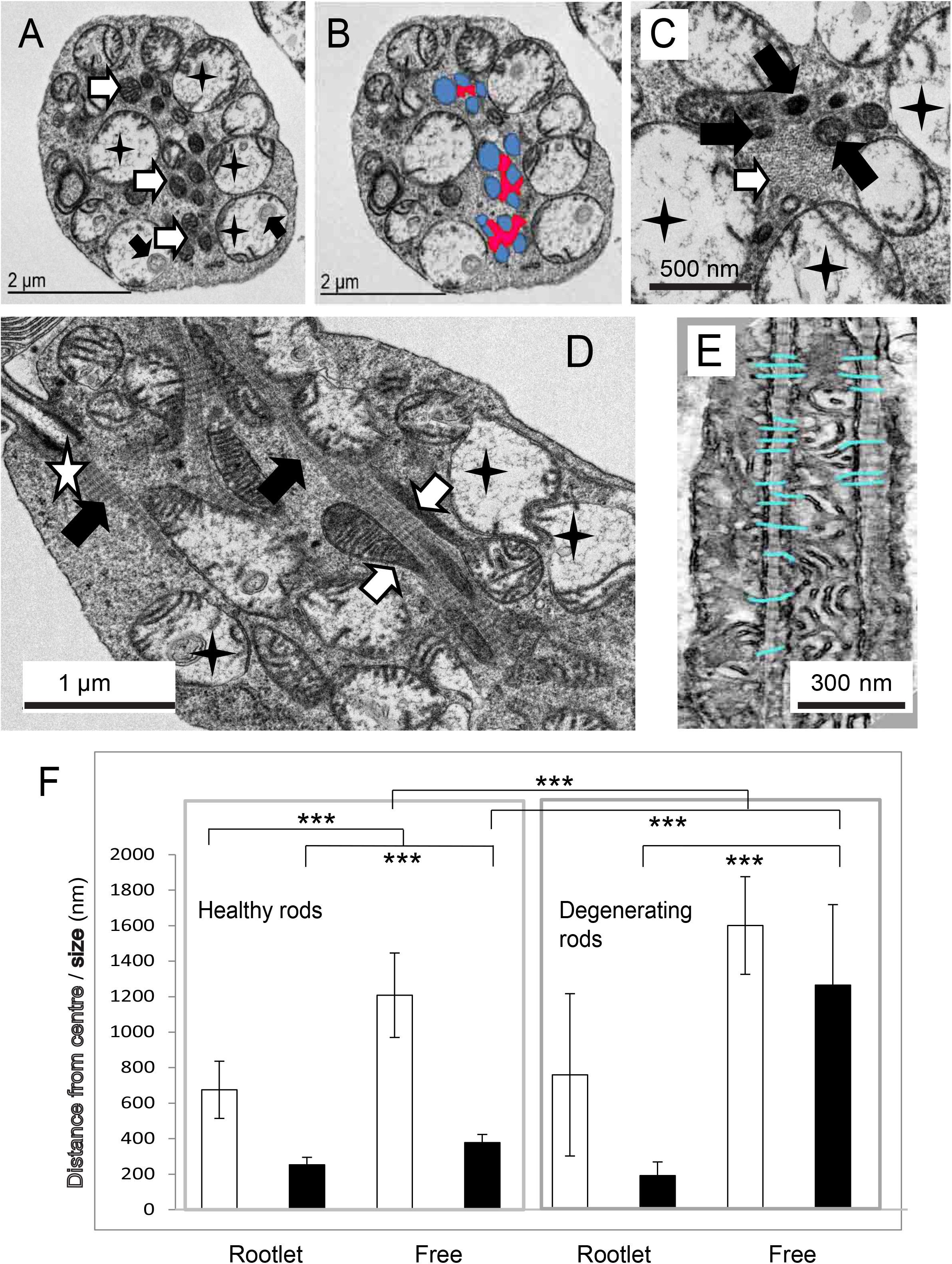
Mitochondria in stressed rods. A: TEM of a transverse section through a stressed rod showing swollen mitochondria with few cristae (black stars). Note the membranous inclusions (black arrows). Narrow, healthy-looking, dense, cristae-rich mitochondria are associated with the rootlet (white arrows). B: C: Close up of the rootlet which appears as a fibrous matrix (white arrow), dense mitochondria (black arrows) swollen mitochondria (black stars). D: Longitudinal section of the rod showing mitochondrial extensions (white arrows) aligned with the rootlet (black arrows). The white star indicates the ciliary basal body. E: Mitochondria cristae aligning with the rootletin repeats on the ciliary rootlet. F: Histogram showing the width (size) and distance from the centre of the photoreceptor of rootlet-associated (Rootlet) and non-rootlet-associated (Free) mitochondria in healthy-looking and stressed rods.

In these rods we observed swollen mitochondria containing membranous whorls. These were less dense than those seen in cones, often comprising just a few invaginations. Neck-like connections to the mitochondrial inter membrane space (MIM) were occasionally visible (Fig 7A). The MIM in this region appeared electron-lucent and depleted, but unlike cones, the mitochondrial membranes in the vicinity of the aggregate are often reduced to a single membrane. In most cases the outer membrane is missing, in others the inner one is absent. Mitochondria with these denuded regions are often visible in the cytoplasm, not only in association with the plasma membrane. When these single-membrane domains contact either the plasma membrane (Fig. 7B-D) or another swollen mitochondrion, we see evidence of a series of membrane fusions resulting in material being ejected from the cell (Fig. 7E) or being exchanged between mitochondria (Fig. 7F).

**Figure 7.**
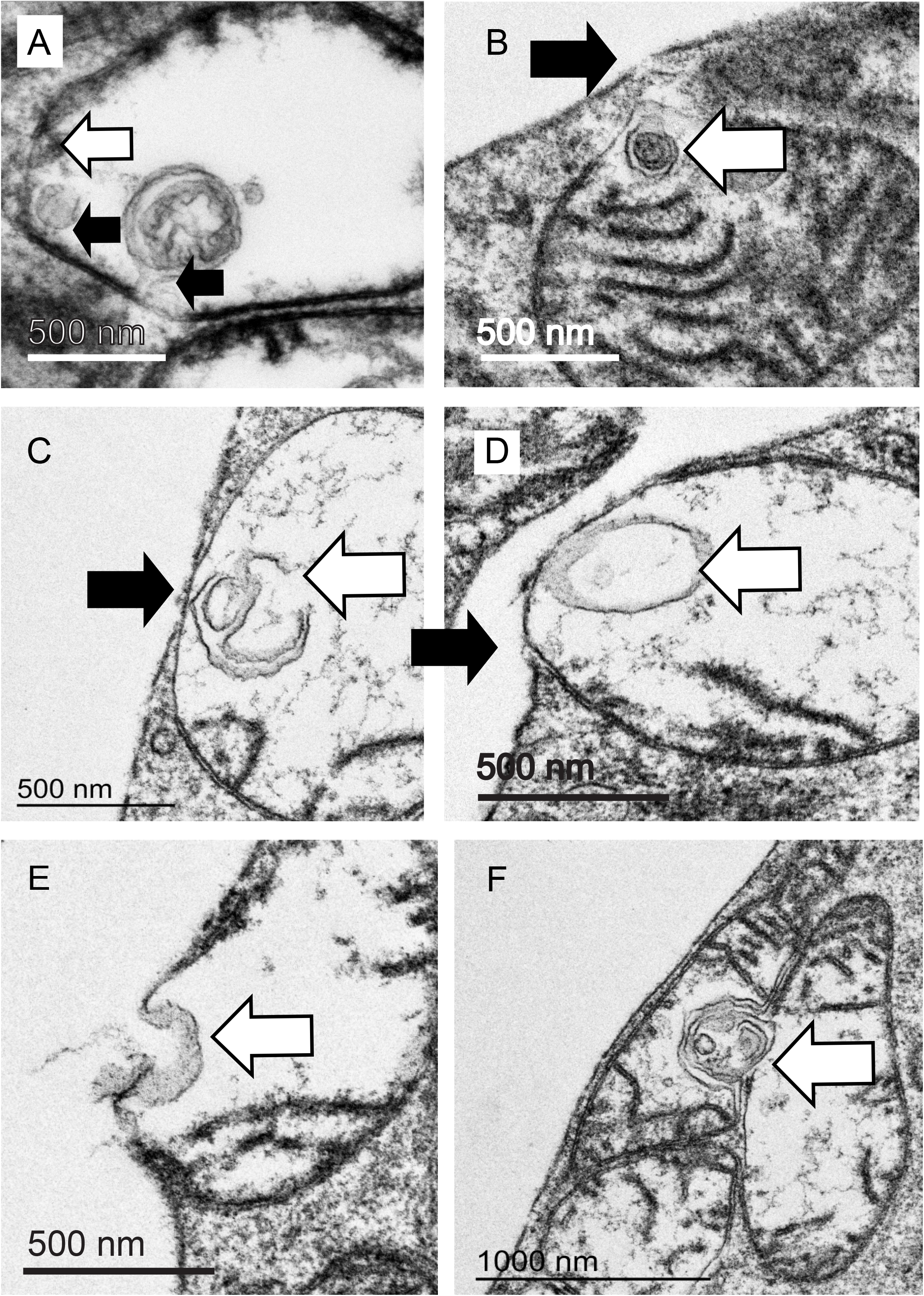
Mitochondria in stressed rods expel and exchange material. A: Swollen mitochondrion in a rod showing few cristae and several internal membranous aggregates connected (presumed) to the inner mitochondrial membrane by a thin ‘neck’(black arrows). Note the mitochondrial outer membrane is ruptured in this domain (white arrow). B-D: The ruptured mitochondrial membrane engaged with the plasma membrane of the photoreceptor (white arrow: membrane aggregate, black arrow: tear in plasma membrane). E: Multiple membrane fusions result in expulsion of contents. F: Fusion may occur between mitochondria resulting in exchange of membranous contents (white arrow) and possible mitochondrial fusion.

In some regions we identified what appear to be patches of more extensive retinal stress where stressed/degenerating rods with polymorphic mitochondria were interspersed between apparently healthy rods and cones (Fig. 8A). The nuclei and outer segments of these rods appeared normal (data not shown), suggesting that the cells were potentially still alive if not functional. In these regions the rod inner-segments contained little cytoplasm and were tightly packed with abnormal mitochondria containing electron-dense membranous aggregates (Fig. 8B). Rod mitochondria in neighbouring, healthy-looking cells were narrow and elongated (Fig. 8C). Only a few such mitochondria remain in some cells, and those we saw were restricted to the core of the cell. In the most extreme cases of degeneration the mitochondria appeared bowl-shaped, were closely opposed to the plasma membrane and had a characteristic open-mouthed appearance from which electron-dense material was seen to spew out of the cell (Fig. 8D-F). Expelled material is fibrous or membranous and of varying density, suggesting it unfolds or disperses as it exits. It hung in the inter-photoreceptor space; presumably associated with the inter-photoreceptor matrix (Fig. 8B).

**Figure 8.**
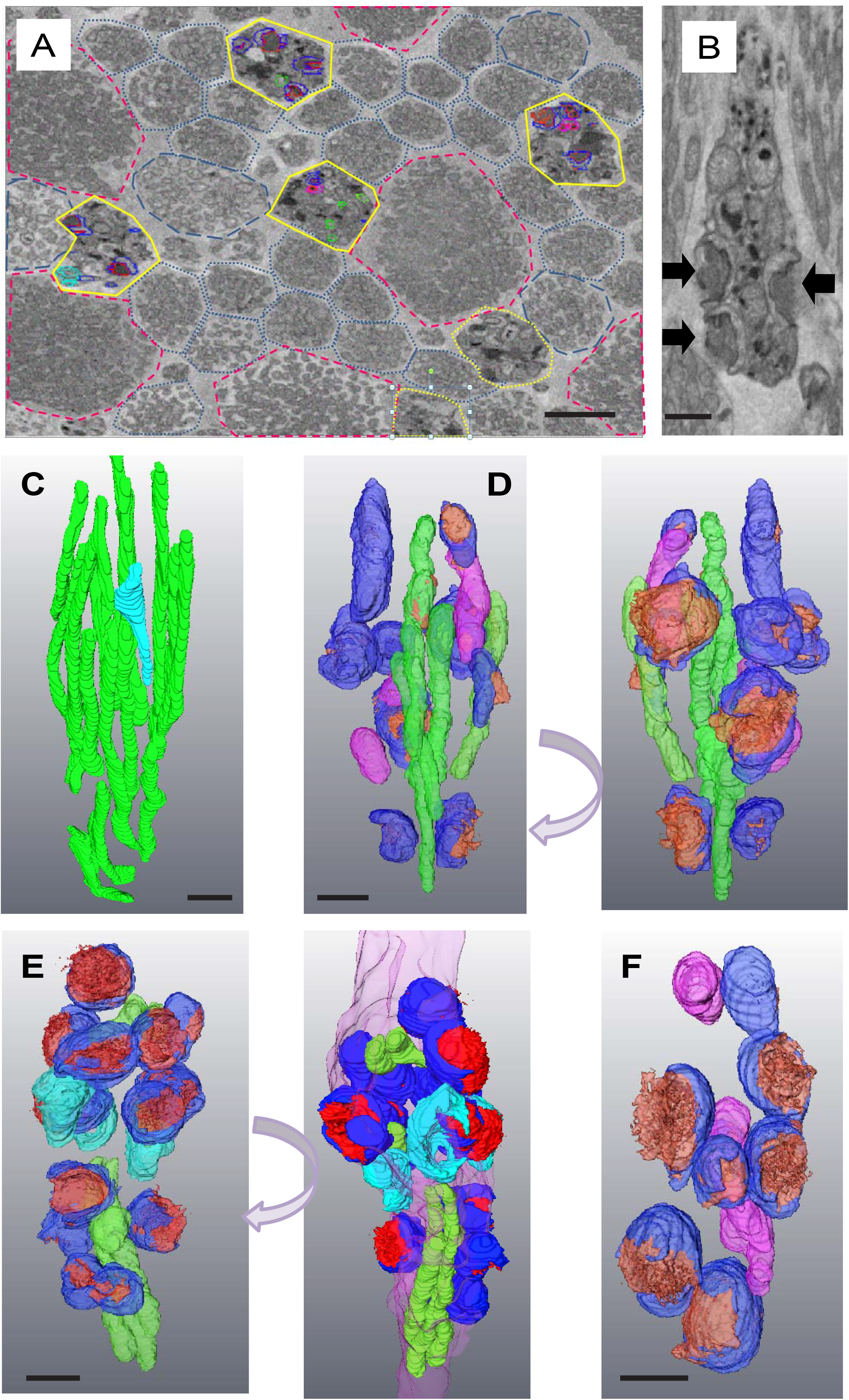
Late stage of degenerating primate rods. A: Low magnification SBF-SEM section of a cluster of late-stage stressed/degenerating rods (high-lighted in yellow). Healthy-looking rods are outlined with blue dots, those containing some swollen mitochondria with blue dashes. Healthy cones are outlined with red dashes. B: Single transverse SBF-SEM section through a late-stage stressed rod. Black arrows indicate electron-dense material being expelled from the mitochondria through the plasma membrane of the photoreceptor. C: Partial segmentation of mitochondria from a healthy rod. The one pale blue mitochondrion shows some swelling. B: Rotations of a rod demonstrating degeneracy. D-F: Rotations 3D reconstructions of segmented mitochondria from rods in various late stages of degeneration. (Green: ‘healthy’ elongated, dense mitochondria, magenta: Swollen mitochondria with electron-lucent matrices, dark blue: mitochondria associated with the plasma membrane expelling electron dense contents (red), cyan: mitochondria associated with the plasma membranes that have ejected all their contents. Scale bars 500 nm

## Discussion

In this paper we describe a potential mechanism by which mitochondrial populations in aging primate photoreceptors preserve healthy mitochondrial domains by linking them to the ciliary rootlet whilst aggregated mitochondrial components are expelled from stressed photoreceptors through the plasma membrane.

Using TEM and SBF-SEM to reveal 3D structures we identified a diverse range of mitochondrial forms in healthy and apparently stressed photoreceptors. In healthy cones we see an apical-distal asymmetry in mitochondrial shape with those most distal, closest to the photoreceptor outer segment, being of larger diameter. This suggests that even within the ellipsoid of a healthy cone the mitochondria are not identical. There may be polarisation of fission-fusion along the apical-distal axis.

In rods and cones cortical mitochondria are closely associated with the plasma membrane and are tubular, elongated and narrow. Away from this zone the inner mitochondria are shorter and a little fatter. The closeness and extent of the association of the cortical mitochondria with the plasma membrane suggests that some kind of contact site is formed between the mitochondrion outer membrane and the plasma membrane. This may be of the form identified in mouse rod photoreceptors ^33^. We identified little evidence of the trans-cellular mitochondrial alignment described in mouse, the photoreceptors being mostly too far apart to allow apposition of such membranes. The photoreceptors are much less tightly packed together in primate retinae than those of rodents in the vicinity of the ellipsoid, with significant inter-photoreceptor matrix between them, making apposition improbable (Supplementary Fig. 1).

We identified thin, tubular subdomains of a sub-population of mitochondria associated with the ciliary rootlets of both rods and cones. These domains have a dense matrix, are free of membranous aggregates and contain abundant, closely-packed cristae. Mitochondria have been shown to associate with rootlets in the photoreceptors of the cow ^31^ and owl monkey ^30^ where there is evidence of correlation between the arrangement of cristae and the rootlet striations, something we also observe here. This is indicative of a cross-link between the rootlet and the mitochondrial contact site and cristae organising system (MICOS). We have seen similar results in rabbit photoreceptors (Supplemental Fig. 2) but not in mice (data not shown).

In mammals mitochondrial fission is regulated by dynamin-related protein (Drp1) and its adaptors Fis1, Mff and Mief1 ^21,24,34^. Drp1 causes fission by encircling the mitochondrion and forming a constriction site as a result of its GFPase activity ^35^. Dynamin-related mitochondrial fission proteins such as Drp1 have evolved to polymerise around mitochondria of a particular diameter ^36^. If, due to swelling, the mitochondrion has too great a circumference, there would be reduced opportunity for fission events.

Mechanical force *per se* is sufficient to recruit the integral mitochondrial fission factor Mff to sites of mitochondrial narrowing ^37,38^. This, in turn, recruits Drp1 and results in successful fission ^39^. Binding to the rootlet presumably facilitates generation of a force which promotes elongation of the mitochondrion into a thin tubule which may be sufficient to recruit Mff and initiate fission. We were able to identify sites along mitochondrial tubules where they were pinched into narrow necks: these may be the sites of fission. We have also noted that in large, healthy cones the core of the photoreceptor, which is where the rootlet is usually to be found, is populated with small, spherical mitochondrial fragments which are likely to be the products of this fission.

In total, this suggests an apical-distal, cortical-core asymmetry of mitochondrial dynamics within single photoreceptor cells in the primate.

Swollen mitochondrial domains with few cristae and individual swollen mitochondria are excluded from the rootlet. We observe invagination of domains of their inner membrane, resulting in formation of electron-dense aggregates in the case of cones, and more diffuse membranous structures in the case of rods. If these aggregates are small, in both rods and cones, they are fully internal to the outer mitochondrial membrane. This excludes the possibility the material is cytoplasmic or lysosomal in origin.

We observe modification or depletion of the mitochondrial outer membrane in the vicinity of electron-dense membranous aggregates which seems to render it susceptible to fusion with the plasma membrane and with other modified domains on other swollen mitochondria. This may result in the release of material from the cell into the inter-photoreceptor space or promote aggregation and fusion of damaged mitochondria. Using 3D electron tomography, we observed what we assume to be highly-stressed rods with very few normal mitochondria, but large swollen mitochondria attached to the plasma membrane in the process of expelling electron-dense contents out of the cell. Any normal mitochondria present were situated in the core of the inner segment; presumably still associated with the rootlet. In cells undergoing mitochondrial remodelling, the nuclei appear normal and the outer-segment is still intact (data not shown). This implies that the cells are not dead and could potentially still be functional. The changes in mitochondrial form and integrity suggest that cytochrome c may have been released, though in some mouse model systems of apoptosis in photoreceptors mitochondrial cytochrome c release does not occur ^40^. We may be observing a very early stage of cell death, or photoreceptors may be refractory to this classical pathway.

Our observations of mitochondrial expulsion of electron-dense aggregates from the photoreceptor are broadly in agreement with those described in zebrafish ^41^. In this elegant paper the authors refer to similar structures they identified in mouse. We do not know if this process is conserved across species; but given that it has been clearly demonstrated in fish, and by us in primates, it seems likely.

Piecemeal mitochondrial degradation, characterised by the production of mitochondrial derived vesicles (MDVs) has been described as a mechanism to isolate damaged material (lipid, protein and mitDNA) from mitochondria and thus purify them ^42,43^. The damaged material has been shown to fuse with a subset of peroxisomes ^44^ or components of the late endosomal pathway such as multivesicular bodies or lysosomes^45^ where it can be disposed of or recycled. MDV production occurs during early stages of ROS-induced damage. If damage progresses to a point at which the mitochondria is completely depolarised, redundant or irreparably damaged, it can also be eradicated by a form of autophagy known as mitophagy ^46^. This is a slower process that is initiated by components that induce MDV formation; but which then recruits the autophagy machine ^47,48^. Conventional piecemeal MDV recycling and mitophagy, superficially at least, appear to be quite distinct from the process we have identified in primate photoreceptors. Our examination of aging or dying photoreceptors in the primate did not reveal the presence of autophagosomes or accumulations of large numbers of multivesicular bodies or lysosomes (our unpublished observations).

Primate rods appear to be more susceptible to photo-damage than cones ^49^ and are lost in an age-dependent manner leaving cones in a less dense photoreceptor matrix, which is not the case in mice where cones die before rods ^50^. Hence, we see far less dramatic changes in mitochondria in cones, though we do see some occasions of mitochondrial tubule formation on the surface of rootlets and can easily identify expulsion of mitochondrial contents from a minority of mitochondria in most cones from older individuals.

This begs the question as to the fate of the purged material which is rich in oxidised mitochondrial lipids and mitochondrial proteins. The mitochondrial debris is ultimately likely to be phagocytosed by the retinal pigmented epithelium ^51^ along with a portion of the outer segment. If this material were generated in significant quantity, as it appears to be in some of the examples we identified, it could overwhelm the RPE or interfere with normal outer-segment phagocytosis and end up being secreted basally, contributing to the basal lateral deposits and drusen (fatty, bleb-like deposits) associated with aging and AMD.

The protein composition of druse provides circumstantial evidence that this might be the case. Drusen are enriched in proteins that have undergone oxidation. Mitochondrial cytochrome c oxidase subunit 5B, mitochondrial aldehyde dehydrogenase and 2-oxoglutarate carrier protein are elevated in basolaminar deposits ^52^ and the mitochondrial inner membrane protein ATP synthase F1 subunit ATP5B is a common and abundant component of drusen ^53^. Cardiolipin, the characteristic phospholipid of mitochondria, has the capacity to initiate the classical complement cascade, remnant deposits of which make up some of the protein contribution to drusen ^54,55^. Cardiolipin also binds to complement factor H (CFH) ^56^, mutations in which are associated with age-related macular degeneration. Mitochondrial detritus expelled from photoreceptors in the atypical manner described, could not only contribute to basolaminar deposits or drusen, but initiate an unproductive complement response. This mechanism provides an alternative source of some of the lipids which make up the bulk of drusen ^57^ and other age-related lipid deposits.

## Authors’ contributions

MJH designed and performed the experiments and wrote the manuscript. DTW assisted in data acquisition and provided discussion. JHK, MBP and GJ provided sample material, valued discussion and edited the manuscript. We would like to thank Professor Tim Levine for comments on the manuscript.

## Acknowledgments

We thank Public Health England for their assistance in obtaining the tissues used. We also thank Sandie Holmes from PHE for her invaluable assistance during tissue collection.

## Figure legends

**Supplementary figure 1.**
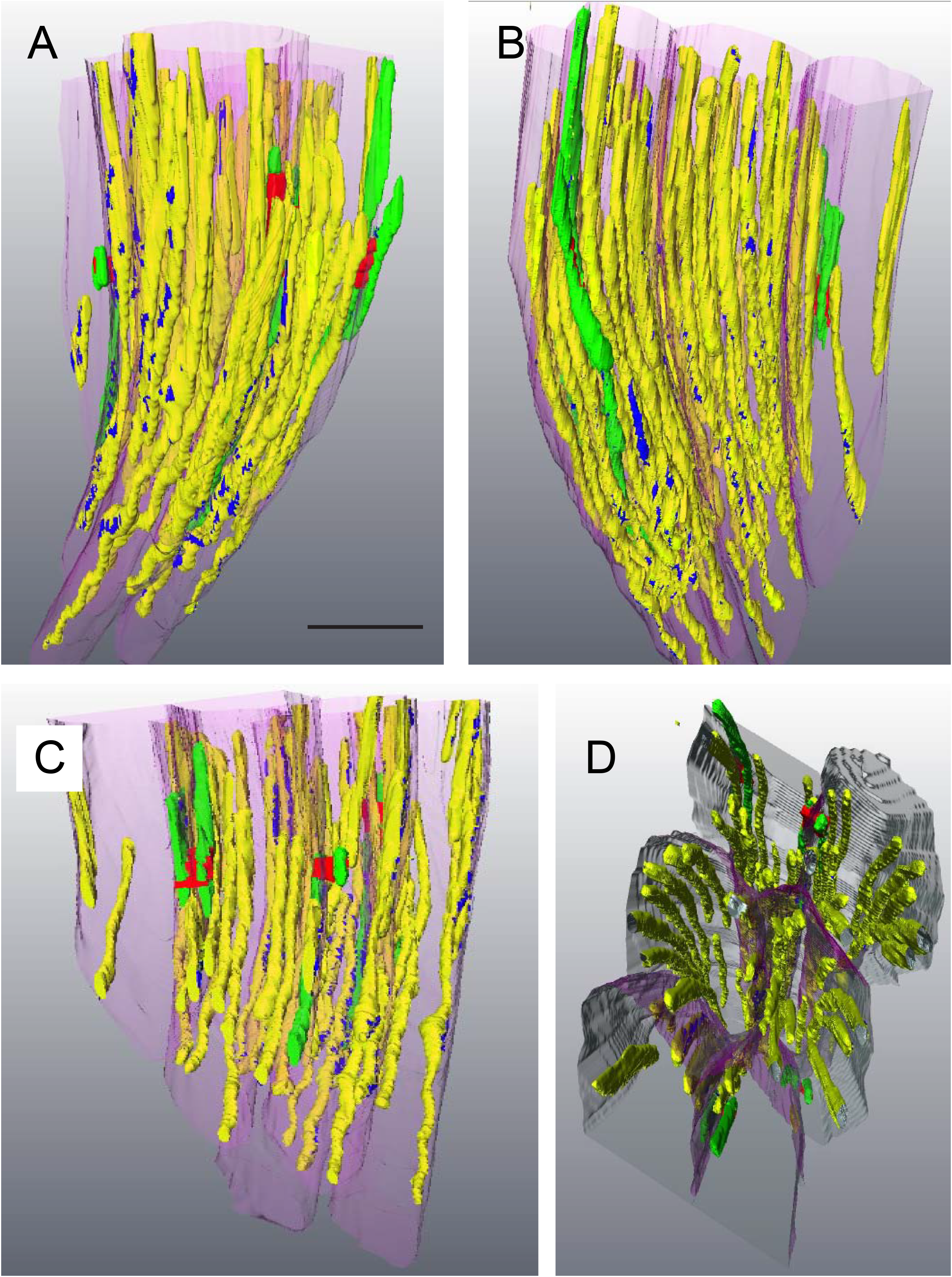
There are few trans-cellular mitochondria contacts between macaque rod photoreceptors. Partial reconstruction of cortical, plasma membrane-associated mitochondria: showing trans-cellular alignment between neighbouring cells (green). Potential contact sites are labelled red. Mitochondria which do not align with others in neighbouring cells are yellow. Their contacts with the plasma membrane are high-lighted in dark blue. A-C: axial rotations of six rod photoreceptors. D: the same group rotated so as to show how these cortical mitochondria are intimately associated with the plasma membrane.

**Supplementary figure 2.**
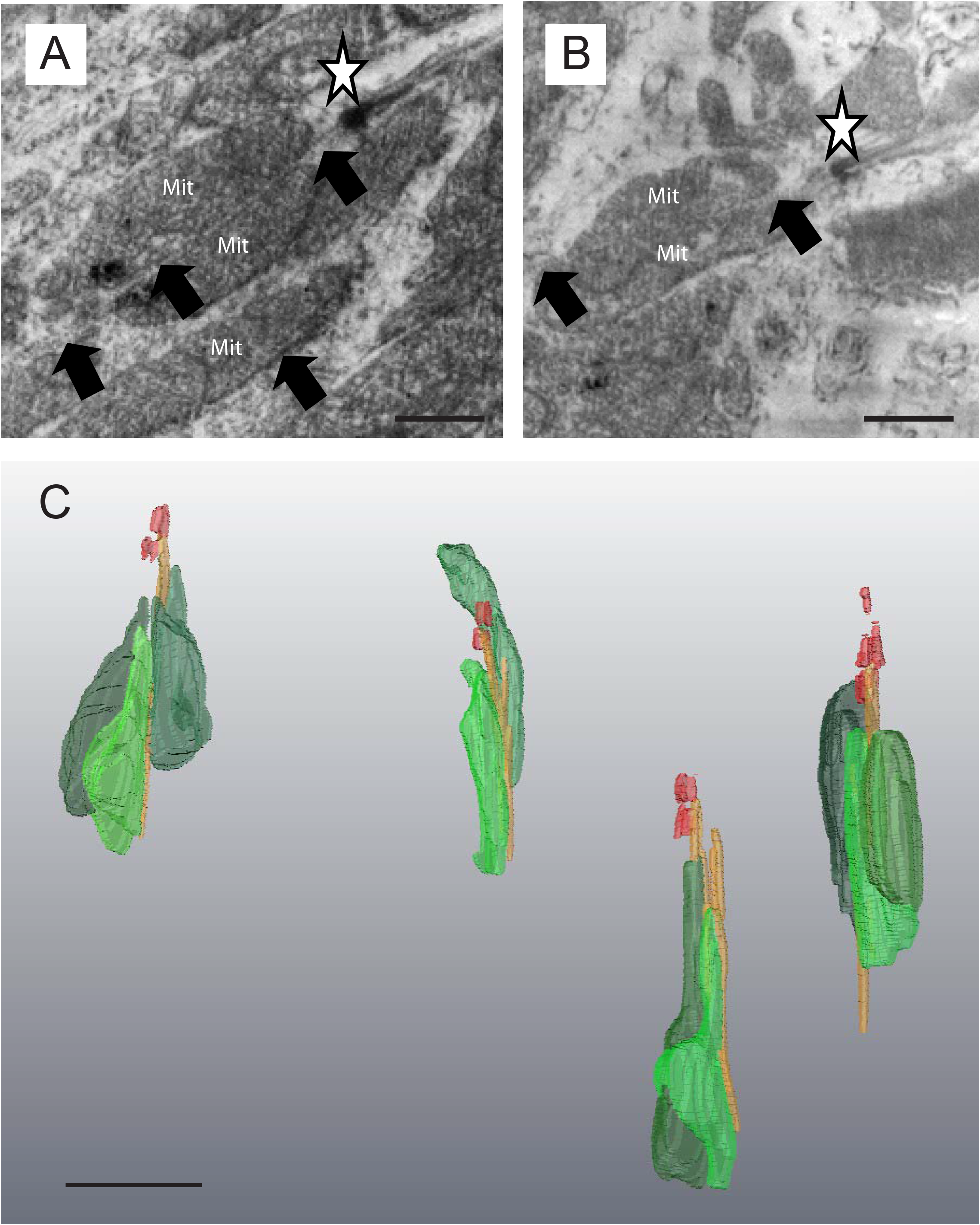
Alignment of mitochondria with the rootlet in rabbit rods. A-B: Single image from a SBF-SEM stack showing mitochondrial association with the ciliary rootlet in the vicinity of the basal body. The ciliary rootlet is just visible as a punctate line running between the mitochondria. Mitochondria (Mit), basal body (white star), rootlet (black arrows). C: 3-dimensional reconstruction from SBF-SEM stacks showing mitochondria associating with the rootlet. Mitochondria (green), paired basal bodies (red), rootlet (orange).

